# An open-source *k*-mer based machine learning tool for fast and accurate subtyping of HIV-1 genomes

**DOI:** 10.1101/362780

**Authors:** Stephen Solis-Reyes, Mariano Avino, Art F.Y. Poon, Lila Kari

## Abstract

For many disease-causing virus species, global diversity is clustered into a taxonomy of subtypes with clinical significance. In particular, the classification of infections among the subtypes of human immunodeficiency virus type 1 (HIV-1) is a routine component of clinical management, and there are now many classification algorithms available for this purpose. Although several of these algorithms are similar in accuracy and speed, the majority are proprietary and require laboratories to transmit HIV-1 sequence data over the network to remote servers. This potentially exposes sensitive patient data to unauthorized access, and makes it impossible to determine how classifications are made and to maintain the data provenance of clinical bioinformatic workflows. We propose an open-source supervised and alignment-free subtyping method (KAMERIS) that operates on *k*-mer frequencies in HIV-1 sequences. We performed a detailed study of the accuracy and performance of subtype classification in comparison to four state-of-the-art programs. Based on our testing data set of manually curated real-world HIV-1 sequences (*n* = 2, 784), Kameris obtained an overall accuracy of 97%, which matches or exceeds all other tested software, with a processing rate of over 1,500 sequences per second. Furthermore, our fully standalone general-purpose software provides key advantages in terms of data security and privacy, transparency and reproducibility. Finally, we show that our method is readily adaptable to subtype classification of other viruses including dengue, influenza A, and hepatitis B and C virus.

## Introduction

Subtype classification is an important and challenging problem in the field of virology. Subtypes (also termed clades or genotypes) are a fundamental unit of virus nomenclature (taxonomy) within a defined species, where each subtype corresponds to a cluster of genetic similarity among isolates from the global population. Defined subtype references for hepatitis C virus, for example, can diverge by as much as 30% of the nucleotide genome sequence [1], but there is no consistent threshold among virus species. Many virus subtypes are clinically significant because of their associations with variation in pathogenesis, rates of disease progression, and susceptibility to drug treatments and vaccines [2]. For example, the HIV-1 subtypes originated early in the history of the global pandemic [3] and have diverged by about 15% of the nucleotide genome sequence [4]. Rates of disease progression vary significantly among HIV-1 subtypes and classifying newly diagnosed infections by their genetic similarity to curated reference subtypes [5] is a recommended component for the clinical management of HIV-1 infection [6, 7]. Consequently, a number of algorithms have been developed for the automated determination of HIV-1 subtypes from genetic sequence data [8–10].

Today, there are important practical considerations that HIV-1 subtyping algorithms should meet. These include:

1. **High Accuracy and Performance:** The cost of sequencing is rapidly decreasing and the amount of sequence data increasing due to next-generation sequencing (NGS) technologies. Thus, in addition to being accurate, software must be computationally fast and scalable in order to handle rapidly growing datasets.
2. **Data Security and Privacy:** Policy, legal, and regulatory issues can prohibit patient sequence data from being transmitted to an external server on the Internet. In addition, concerns around privacy policies and the possibility of data breaches can cause issues for researchers and clinicians. For these reasons, software should be made available in an offline, standalone version.
3. **Transparency:** With closed-source or proprietary software, it can be impossible to determine precisely how classification determinations are made. An open-source implementation gives full visibility into all aspects of the classification process.
4. **Reproducibility:** Relying on an externally-hosted service can make it impossible to determine which version of the software has been used to generate subtype classifications. This makes it difficult to guarantee that classification results can be reproduced, and reproducibility is generally recognized as a necessary component of clinical practice.

In our effort to develop a general sequence classification method satisfying the above considerations, we propose a simple, intuitive, general-purpose, highly-efficient technique based on *k*-mer frequency vectors for supervised nucleotide sequence classification, and we release an open-source software implementation of this method (designated KAMERIS) under a permissive open-source license.

### Alignment-free subtyping

Most subtype classification methods for HIV-1 require the alignment of the input sequence against a set of predefined subtype reference sequences [11], which enables the algorithm to compare homologous sequence features [12–14]. For example, the NCBI genotyping tool [15] computes BLAST similarity scores against the reference set for sliding windows along the query sequence. Other methods such as REGA [9] and SCUEAL [10] reconstruct maximum likelihood phylogenies from the aligned sequences: REGA (version 3.0) reconstructs trees from sliding windows of 400bp from the sequence alignment and quantifies the confidence in placement of the query sequence within subtypes by bootstrap sampling (bootscanning) [16]. Alignment-based methods are relatively computationally expensive, especially for long sequences; the heuristic methods require a number of *ad hoc* settings, such as the penalty for opening a gap; and alignment method may not perform well on highly-divergent regions of the genome. To address these limitations, various alignment-free classification methods have been proposed. Some of them make use of nucleotide correlations [17], or sequence composition (*e.g.* COMET [8] and [18]). Other methods include those based on restriction enzyme site distributions, applied to the subtyping of human papillomavirus (HPV), hepatitis B virus (HBV) and HIV-1 (CASTOR [19]); based on the “natural vector” which contains information on the number and distribution of nucleotides in the sequence, applied to the classification of single-segmented [18] and multi-segmented [20] whole viral genomes, as well as viral proteomes [21]; based on neural networks using digital signal processing techniques to yield “genomic cepstral coefficient” features, applied to distinguishing four different pathogenic viruses [22]; based on different genomic materials (namely DNA sequences, protein sequences, and functional domains), applied to the classification of some viral species at the order, family, and genus levels [23]; and based on interpolated Markov models, applied to the phylogenetic classification of metagenomic samples [24].

### *k*-mer-based classifiers

The use of *k*-mer (substrings of length *k*) frequencies for phylogenetic applications started with Blaisdell, who reported success in constructing accurate phylogenetic trees from several coding and non-coding nDNA sequences [25] and some mammalian alpha and beta-globin genes [26]. Other authors [27–31] have observed that the excess and scarcity of specific *k*-mers, across a variety of different DNA sequence types (including viral DNA in [27]), can be explained by factors such as physical DNA/RNA structure, mutational events, and some prokaryotic and eukaryotic repair and correction systems. Typically, *k*-mer frequency vectors are paired together with a distance function in order to measure the quantitative similarity between any pair of sequences. Studies measuring quantitative similarity between DNA sequences from different sources have been performed, for instance using the Manhattan distance [32, 33], the weighted or standardized Euclidean distance [34, 35], and the Jensen-Shannon distance [36, 37]. Applications of these distances and others have been compared and benchmarked in [38–41], and detailed reviews of the literature can be found in [42–45].

In the context of viral phylogenetics, *k*-mer frequency vectors paired with a distance metric have been used to construct pairwise distance matrices and derive phylogenetic trees, *e.g.*, dsDNA eukaryotic viruses [46], or fragments from Flaviviridae genomes [47]. Other studies have investigated the multifractal properties of *k*-mer frequency patterns in HIV-1 genomes [48], and the changes in dinucleotide frequencies in the HIV genome across different years [49]. We used *k*-mer frequency vectors to train supervised classification algorithms. Similar approaches have previously been explored (with different classifiers than those used here), for example to subtype Influenza and classify Polyoma and Rhinovirus fragments [50], to predict HPV genotypes [51, 52], to classify whole bacterial genomes to their corresponding taxonomic groups at different levels [53], to classify whole eukaryotic mitochondrial genomes [54–57], to classify microbial metagenomic samples [58], to predict virus-host relationships for some bacterial genera [59], and to identify viral sequences in metagenomic samples [60].

To evaluate our method, we curated manually-validated testing sets of ‘real-world’ HIV-1 data sets. We assessed fifteen classification algorithms and conclude that for these data the SVM-based classifiers, multilayer perceptron, and logistic regression achieved the highest accuracy, with the SVM-based classifiers also achieving the lowest running time out of those. We measured classification accuracy and running time for *k*-mers of length *k* = 1 … 10 and found that *k* = 6 provides the optimal balance of accuracy and speed. Overall, our open-source method obtains a classification accuracy average of 97%, with individual accuracies equal or exceeding other subtyping methods for most datasets, and processes over 1,500 sequences per second. Our method is also applicable to other virus datasets without modification: we demonstrate classification accuracies of over 90% in all cases for full-length genome data sets of dengue, hepatitis B, hepatitis C, and influenza A viruses.

## Methods

### Supervised classification

First, we needed to determine which supervised classification method would be the most effective for classifying virus sequences, using their respective *k*-mer frequencies as feature vectors (numerical representations). We trained each of 15 classifiers (Table 2) on a set *S* = {*s*_1_, *s*_2_, … *s* _*n*_} of nucleotide sequences partitioned into groups *g*_1_, *g*_2_, …, *g*_*p*_. Given as input any new, previously unseen, sequence (*i.e.*, not in the dataset *S*), the method outputs a prediction of the group *g*_i_ that the sequence belongs to, having ‘learned’ from the training set *S* the correspondence between the *k*-mer frequencies of training sequences and their groups. The feature vector *F*_*k*_(*s*) for an input sequence *s* was constructed from the number of occurrences of all 4^*k*^ possible *k*-mers (given the nucleotide alphabet {*A*, *C*, *G*, *T*}), divided by the total length of *s*. Any ambiguous nucleotide codes (*e.g.*, ‘*N*’ for completely ambiguous nucleotides) were removed from *s* before computing *F*_*k*_(*s*).

**Table 1.** Statistics for the manually curated testing datasets. The first author, year, and reference number for the publication associated with each data set is listed under the ‘Source’ column heading. The historically most prevalent HIV-1 subtype(s) is indicated under the ‘Subtype’ column heading.

| Source | Country | Subtype | Count | Sequence length (nt) |  |  |
| --- | --- | --- | --- | --- | --- | --- |
|  |  |  |  | Average | Min. | Max. |
| Nadai (2009) [86] | Haiti | B | 66 | 1024.0 | 1024 | 1025 |
| Niculescu (2015) [87] | Romania | F | 97 | 1301.2 | 1257 | 1302 |
| Paraschiv (2017) [88] | Romania | F | 86 | 1295.9 | 1164 | 1299 |
| Rhee (2017) [89] | Thailand | CRF01_AE | 282 | 703.8 | 633 | 756 |
| Sukasem (2007) [90] | Thailand | CRF01_AE | 221 | 286.4 | 270 | 288 |
| Eshleman (2001) [91] | Uganda | A/D | 102 | 1261.2 | 1260 | 1302 |
| Ssemwanga (2012) [92] | Uganda | A/D | 72 | 1025.0 | 1025 | 1025 |
| Wolf (2017) [93] | USA | B | 1653 | 1020.8 | 868 | 1080 |
| TenoRes Study Group (2016) [94] | South Africa | C | 102 | 1001.4 | 921 | 1209 |
| van Zyl (2017) [95] | South Africa | C | 59 | 1056.7 | 1002 | 1070 |
| Huang (2003) [96] | Reference panel | n/a | 44 | 1189.9 | 1187 | 1190 |
| **Overall** |  |  | **2784** | **960.4** | **270** | **1302** |

**Table 2.** Classification accuracy scores after 10-fold cross-validation and running times averaged over five runs for each of the fifteen classifiers at *k* = 6, for the full set of 6625 whole HIV-1 genomes from the LANL database.

| Classifier | Accuracy | Running time |  |
| --- | --- | --- | --- |
|  |  | Mean | Standard deviation |
| cubic-svm | 96.66% | 59.7s | 0.81s |
| quadratic-svm | 96.59% | 58.3s | 0.91s |
| linear-svm | 96.49% | 57.7s | 1.60s |
| multilayer-perceptron | 95.49% | 60.6s | 0.94s |
| logistic-regression | 95.32% | 102.0s | 0.92s |
| 10-nearest-neighbors | 93.97% | 44.3s | 0.68s |
| nearest-centroid-median | 93.95% | 34.0s | 0.83s |
| nearest-centroid-mean | 93.84% | 33.7s | 0.33s |
| decision-tree | 93.53% | 62.3s | 1.03s |
| random-forest | 93.07% | 43.7s | 0.77s |
| sgd | 91.10% | 37.4s | 1.32s |
| gaussian-naive-bayes | 87.75% | 34.0s | 0.80s |
| lda | 77.76% | 36.0s | 1.37s |
| qda | 75.13% | 38.3s | 0.48s |
| adaboost | 64.85% | 159.3s | 1.21s |

Next, we processed the feature vectors for more efficient use by classifiers. We rescaled the *k*-mer frequencies in *F*_*k*_(*s*) to have a standard deviation of 1, which satisfies some statistical assumptions invoked by several classification methods. In addition, we performed dimensionality reduction using truncated singular value decomposition [61] to reduce the vectors to 10% of the average number of non-zero entries of the feature vectors. This greatly reduces running time for most classifiers while having a negligible effect on classification accuracy.

Finally, we trained a supervised classifier on the vectors *F*_*k*_(*s*). Supervised classifiers, in general, can be intuitively thought of as constructing a mapping from the input feature space to another space which in some sense effectively separates each training class. As a concrete example, the support vector machine (SVM) classifier maps the input space to another space of equal or higher dimensionality using a kernel function, and then selects hyperplanes that represent the largest separation between every pair of classes. Those hyperplanes induce a partition on the transformed space which is then used for the classification of new items. We tested fifteen different specific classifier algorithms: 10-nearest-neighbors [62] with Euclidean metric and uniform weights (`10-nearest-neighbors`); nearest centroid, to class mean (`nearest-centroid-mean`) and to class median (`nearest-centroid-median`) [63]; logistic regression with L2 regularization, one-vs-rest as the multiclass generalization, stopping tolerance of 10^−4^, and regularization strength (λ) of 1 (`logistic-regression`) [64]; SVM with the linear (`linear-svm`), quadratic (`quadratic-svm`), and cubic (`cubic-svm`) kernel functions, with penalty parameter of 1 and stopping tolerance of 10^−3^ [65]; SVM with stochastic gradient descent learning (maximum 5 training epochs) and linear kernel function (sgd) [66]; decision tree with Gini impurity metric (`decision-tree`) [67]; random forest using 10 decision trees with Gini impurity metric as sub-estimators (`random-forest`) [68]; AdaBoost with maximum 50 decision trees as the weak learners and the SAMME.R real boosting algorithm (`adaboost`) [69, 70]; Gaussian naïve Bayes (`gaussian-naive-bayes`) [71]; linear (`lda`) and quadratic (`qda`) discriminant analysis [72]; and multi-layer perceptron with a single 100-neuron hidden layer, rectified linear unit (ReLU) activation function, the Adam stochastic gradient-based weight optimizer, L2 penalty term of 10^−4^, learning rate of 10^−3^, and maximum 200 epochs (`multilayer-perceptron`) [73, 74]. We used the implementations of these classifiers in the Python library `scikit-learn` [75] – all settings and hyperparameters were left as the defaults given in the library.

For some of the results that follow, we required a method for measuring classification accuracy without the need for a separate testing dataset. To do so, we used 10-fold cross-validation, a technique widely used for assessing the performance of supervised classifiers [76]. *N*-fold cross-validation is performed by taking the given dataset and randomly partitioning it into *N* groups of equal size. Taking each group in turn, we trained a classifier on the sequences outside of the selected group, and then computed its accuracy from predicting the classes of the sequences in the selected group. The outcome of the cross-validation are *N* accuracy values for the *N* distinct, independent training and testing runs. We report the arithmetic mean of those accuracies as the final accuracy measure.

### Unsupervised visualization

Supervised classification requires, by definition, a training set consisting of examples of classes determined *a priori*. However, one may wish to explore a dataset where the groups are not necessarily all known. For the problem of virus subtyping for example, one may suspect the existence of a novel subtype or recombinant. To this end, unsupervised data exploration techniques are useful, and herein we also explore the use of Molecular Distance Maps (MoDMaps), previously described in [41, 77, 78], for this purpose. After computing the vectors *F*_*k*_(*s*), this method proceeds by first constructing a pairwise distance matrix. In this paper, we use the well-known Manhattan distance [79], defined between two vectors *A* = (*a*_1_, … *a*_*n*_) and *B* = (*b*_1_, … *b*_*n*_) as being:

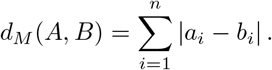

Next, the distance matrix is visualized by classical MultiDimensional Scaling (MDS) [80]. MDS takes as input a pairwise distance matrix and produces as output a 2D or 3D plot, called a MoDMap [81], wherein each point represents a different sequence, and the distances between points approximate the distances from the input distance matrix. As MoDMaps are constrained to two or three dimensions, it is in general not possible for the distances in the 2D or 3D plot to match exactly the distances in the distance matrix, but MDS attempts to make the difference as small as possible.

### Implementation

We have developed a software package called KAMERIS which implements our method. It can be obtained from https://github.com/stephensolis/kameris, and may be used on Windows, macOS, and Linux. KAMERIS is implemented in Python, with the feature vector computation parts implemented in C++ for performance. It is packaged so as to have no external dependencies, and thus is easy to run. The package has three different modes: first, it can train one or more classifiers on a dataset and evaluate cross-validation performance; second, it can summarize training jobs, computing summary statistics and generating MDS plots; and third, it can classify new sequences on already-trained models. More information, including usage and setup instructions, can be found at https://github.com/stephensolis/kameris. All running time benchmarks of our software were performed on an Amazon Web Services (AWS) r4.8xlarge instance with 16 physical cores (32 threads) of a 2.3GHz Intel Xeon E5-2686 v4 processor. We also note that many of the implementations of the classifier algorithms we use are single-threaded and that performance can almost certainly be substantially improved by using parallelized implementations.

### Datasets

In this paper, a variety of different datasets were used to validate the performance of the method. Straightforward reproducibility of results was a priority in the design of this study, and to that end, every sequence and its metadata from every dataset referenced here can be retrieved from our GitHub repository at https://github.com/stephensolis/kameris-experiments. Further, instructions for using KAMERIS to replicate the experiment results are available in S1 Appendix.

In some cases, these datasets had few examples for some classes. Training on classes with very few examples would unfairly lower accuracy since the classifier does not have enough information to learn, so we wish to omit such classes from our analysis. However, the minimum number of examples per class to achieve proper training of a classifier is difficult to estimate; this number is known to be dependent on both the complexity of the feature vectors and characteristics of the classifier algorithm being used [82, 83]. Since we vary both *k* and the classifier algorithms in this study, this makes it especially challenging to determine an adequate minimum class size. Here, we arbitrarily selected 18 as our minimum, so we omitted from analysis any subtype with fewer than 18 sequences. It may be that specific values of *k* and some classifier algorithms work well in scenarios with very small datasets, and we leave this as an open question.

#### Primary dataset

The primary dataset used was the full set of HIV-1 genomes available from the Los Alamos (LANL) sequence database, accessible at https://www.hiv.lanl.gov/components/sequence/HIV/search/search.html. In this database, the option exists of using full or partial sequences – in our analysis, we consider both full genomes and just the coding sequences of the *pol* gene. For the set of whole genomes, the query parameters “virus: HIV-1, genomic region: complete genome, excluding problematic” were used; this gave a total of 6625 sequences with an average length of 8970 bp. For the set of *pol* genes, the query parameters “virus: HIV-1, genomic region: Pol CDS, excluding problematic” were used; this gave a total of 9270 sequences with an average length of 3006 bp. In both cases, the query was performed on May 18, 2017, and at the time, the LANL database reported a last update on May 6, 2017. After removing small classes (see preceding section), this dataset contained 25 subtypes and circulating recombinant forms (CRFs) for the set of whole genomes, and 26 for the set of *pol* genes. The list of subtypes for this dataset and all other datasets described here are available in S2 Appendix. This dataset was used to determine the best value of *k*, the best classifier algorithm, to compare the performance of whole genomes with *pol* gene sequences only, and to produce the MoDMaps of HIV-1. In those experiments, cross-validation was used to randomly draw training and testing sets from the dataset.

#### Evaluation datasets

To evaluate classifiers trained on HIV-1 sequences and subtype annotations curated by the LANL database, we needed testing sets but wanted to avoid selecting them from the same database. We manually searched the GenBank database for large datasets comprising HIV-1 *pol* sequences collected from a region with known history of a predominant subtype, and evaluated the associated publications to verify the characteristics of the study population (Table 1). After selection of the datasets, we wished to obtain labels without relying on another subtyping method. To do so, first we made use of the known geographic distribution of HIV-1 subtypes, where specific regions are predominantly affected by one or two particular subtypes or circulating recombinant forms due to historical ‘founding’ events [84]. Next, we screened each dataset using a manual phylogenetic subtyping process to verify subtype assignments against the standard reference sequences. This was done, essentially, by reconstructing phylogenetic trees to identify possible subtype clusters. A cluster was identified as a certain subtype if it included a specific subtype reference sequence we had initially provided in our datasets. Thus, the first step was to download the most recent set of subtypes reference sequences for the HIV-1 *pol* gene at the LANL database, accessible at https://www.hiv.lanl.gov/content/sequence/NEWALIGN/align.html [85].

We loaded the resulting FASTA file in the eleven datasets from Table 1. We then aligned the datasets with MUSCLE v3.8.425 [97], implemented in AliView 1.19-beta-3 [98], where we also visually inspected the alignments. To avoid overfitting, we searched for the nucleotide model of substitution that was best supported by each dataset using the Akaike Information Criterion (AIC) in jModeltest v2.1.10 [99]. For the dataset US.Wolf2017, the large number of sequences precluded this model selection process, so we chose a General Time Reversible model incorporating an invariant sites category and a gamma distribution to model rate variation among the remaining sites (GTR+I+G); this parameter-rich model is often supported by large HIV-1 data sets, and was similar to the model selected by the authors in the original study [93]. Phylogenetic trees were reconstructed by maximum likelihood using PHYML v20160207 [100] with a related bootstrap support analysis. The resulting trees were visualized and their relative sequences were manually annotated in FigTree v1.4.3 [101].

In order to benchmark performance on this manually curated testing dataset, we required a separate training dataset. Since the subtype annotations from the full set of HIV-1 genomes in the LANL database are typically given by individual authors using unknown methods, they may be incorrect at times, potentially negatively impacting classification performance. Thus, we trained our classifier on the subset of HIV-1 *pol* sequences from the 2010 Web alignment from the LANL database, accessible at https://www.hiv.lanl.gov/content/sequence/NEWALIGN/align.html. This Web alignment dataset is a more curated set of *pol* sequences, and is more likely to be correctly annotated. Specifically, we selected ‘Web’ as the alignment type, ‘HIV1/SIVcpz’ as organism, ‘POL’ as ‘Pre-defined region of the genome’ under ‘Region’, ‘All’ as subtype, ‘DNA’, and ‘2010’ as the year. Any Simian Immunodeficiency Virus (SIV) sequences were manually removed from the query results. This gave a total of 1979 sequences, containing 15 subtypes or CRFs after removal of small classes.

#### Synthetic data

For another experiment, we generated a set of synthetic HIV-1 sequences by simulating the molecular evolution of sequences derived from the curated HIV-1 subtype references. To do so, we used a modified version of the program INDELible [102], assigning one of the subtype reference sequences to the root of a ‘star’ phylogeny with unit branch lengths and 100 tips. The codon substitution model parameters, including the transition-transversion bias parameter and the two-parameter gamma distribution for rate variation among sites, were calibrated by fitting the same type of model to actual HIV-1 sequence data [103]. We adjusted the ‘treelength’ simulation parameter in the INDELIBLE program control file to modify the average divergence between sequences at the tips.

#### Other virus datasets

Finally, we performed experiments with dengue, influenza A, hepatitis B, and hepatitis C virus sequences. The dengue and influenza sequences were retrieved from the National Center for Biotechnology Information (NCBI) Virus Variation sequence database on August 10, 2017. The dengue virus sequences were accessed from https://www.ncbi.nlm.nih.gov/genomes/VirusVariation/Database/nph-select.cgi?taxid=12637 with the query options “Nucleotide”, “Full-length sequences only”, and “Collapse identical sequences” for a total of 4893 sequences with an average length of 10585 bp. Influenza sequences were accessed from https://www.ncbi.nlm.nih.gov/genomes/FLU/Database/nph-select.cgi?go=genomeset with the query options “Genome sets: Complete only”, and “Type: A” for a total of 38215 sequences with an average length of 13455 bp. Hepatitis B sequences were retrieved from the Hepatitis B Virus Database operated by the Institut de Biologie et Chimie des Proteines (IBCP), accessible at https://hbvdb.ibcp.fr/HBVdb/HBVdbDataset?seqtype=0, on August 10, 2017 for a total of 5841 sequences with an average length of 3201 bp. Finally, hepatitis C sequences were retrieved from the Los Alamos (LANL) sequence database, accessible at https://hcv.lanl.gov/components/sequence/HCV/search/searchi.html, on August 10, 2017, using the query options “Excluding recombinants”, “Excluding ‘no genotype”’, “Genomic region: complete genome”, and “Excluding problematic” for a total of 1923 sequences with an average length of 9140 bp. After removal of small classes, our data comprised 4 subtypes of dengue virus, 12 subtypes of hepatitis B, 6 subtypes of hepatitis C, and 56 subtypes of influenza type A.

## Results

Our subtype classification method has two main parameters that may be varied: namely, the specific classifier to be used, and the value *k* of the length of the *k*-mers to count when producing feature vectors. We begin with the full set of full-length HIV-1 genomes from the LANL database, and we perform a separate 10-fold cross-validation experiment for each of the fifteen classifiers listed in the Methods section, and all values of *k* from 1 to 10, that is, 160 independent experiments in total. For each value of *k*, we plot the highest accuracy obtained by any classifier as well as the average running time over the classifiers, see Fig 1. We observe that *k* = 6 achieves a good balance between classifier performance and accuracy, so at *k* = 6, we list the accuracy obtained by each classifier and its corresponding running time averaged over five runs, see Table 2. As can be seen, the SVM-based classifiers, multilayer perceptron, and logistic regression achieve the highest accuracy, with the SVM-based classifiers achieving also the lowest running time out of those.

**Fig 1.**
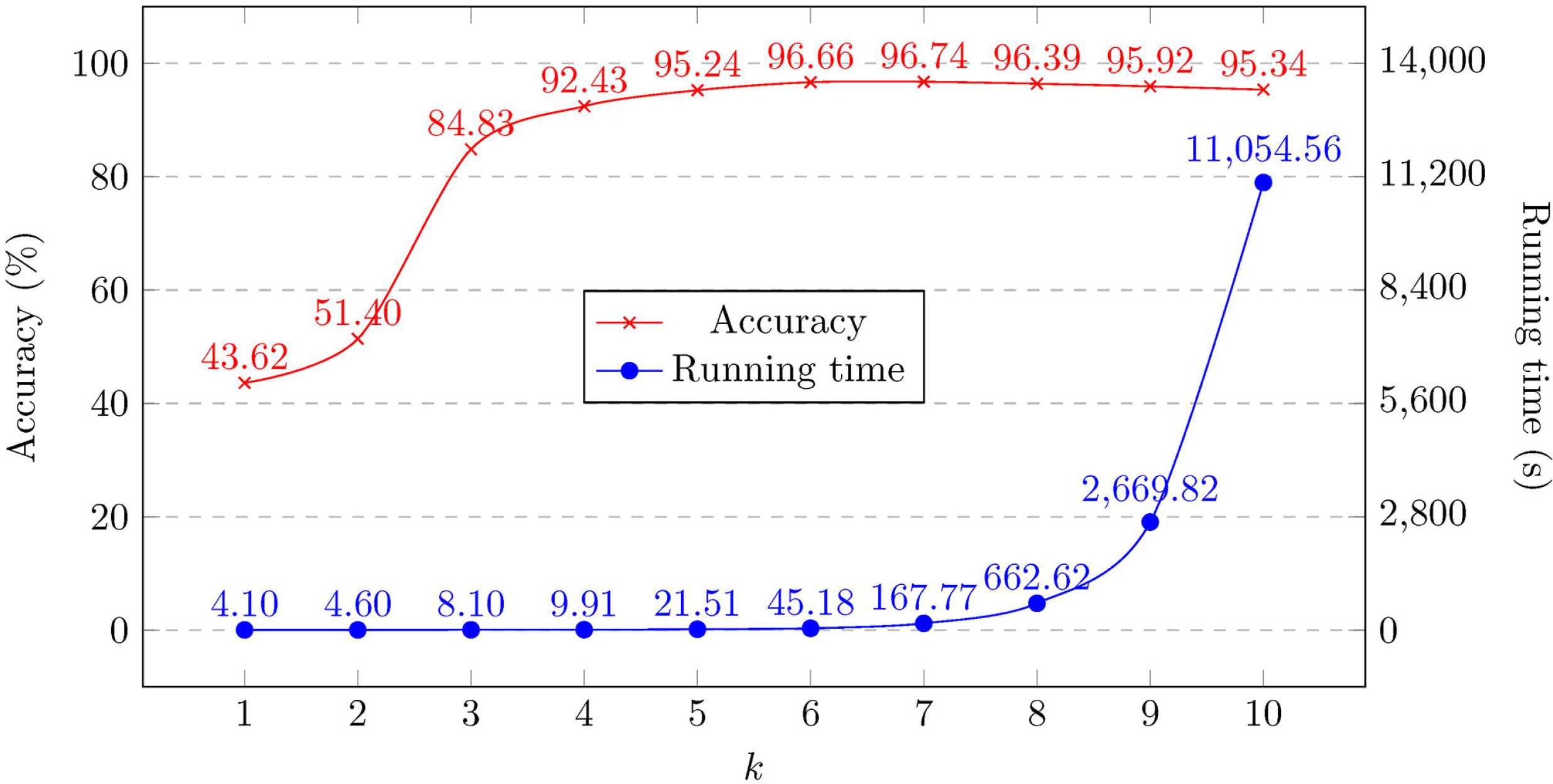
Highest accuracy score and running time averaged across all fifteen classifier algorithms, at different values of *k*, for the full set of 6625 whole HIV-1 genomes from the LANL database.

Since it is typical to have only partial genome sequences available, we repeat the same 10-fold cross-validation at *k* = 6, with the linear SVM classifier, this time with the set of all *pol* genes from the LANL database. We find that the accuracy changes from 96.49% (full-length genomes) to 95.68% (*pol* gene sequences), indicating that the use of partial genomes does not substantially reduce classification performance. Further, we expect that the inclusion of recombinant forms should lower accuracy, since it requires the classifier to accurately distinguish them from their constituent ‘pure’ subtypes. To test this, we repeat the same 10-fold cross-validation at *k* = 6 and with the linear SVM classifier, with the set of all full-length genomes from the LANL database, this time omitting the 17 classes of recombinant forms and leaving only the 9 classes of pure subtypes. We find that the accuracy increases from 96.49% (including recombinants) to 99.64% (omitting recombiants), and in fact only 3 sequences are misclassified in the latter case.

The sequences present in the LANL database are curated to be representative of global HIV-1 diversity, and therefore high classification accuracies on that dataset are, to some extent, to be expected. In order to perform a more challenging benchmark on our algorithm, we compute its accuracy on the eleven selected testing datasets of *pol* gene fragments from Table 1, after training with the set of whole *pol* genes from the LANL 2010 web alignment. Based on the previous performance measurements, we use the linear SVM classifier and *k* = 6. We also perform the same accuracy measurement with four other state-of-the-art HIV subtyping tools: CASTOR, COMET, SCUEAL, and REGA, and show the results in Table 3. In sum, our method (KAMERIS) comes within a few percent of the best tools in all cases, and has the highest average accuracy (both unweighted, and weighted by the number of sequences in each set).

**Table 3.** Classification accuracies for all tested HIV-1 subtyping tools, for each testing dataset; average accuracy both with and without weighting datasets by the number of sequences they contain.

| Source | KAMERIS | COMET | CASTOR | SCUEAL | REGA |
| --- | --- | --- | --- | --- | --- |
| Nadai (2009) [86] | 100.0% | 100.0% | 81.8% | 92.4% | 86.4% |
| Niculescu (2015) [87] | 95.9% | 96.9% | 75.3% | 94.8% | 100.0% |
| Paraschiv (2017) [88] | 91.9% | 73.3%<sup>1</sup> | 46.5% | 68.6% | 87.2% |
| Rhee (2017) [89] | 94.0% | 95.4% | 0.4% | 75.9% | 12.8%<sup>2</sup> |
| Sukasem (2007) [90] | 90.0% | 91.0% | 0.9% | 64.3% | 8.1%<sup>2</sup> |
| Eshleman (2001) [91] | 88.5% | 90.6% | 4.2% | 84.4% | 90.6% |
| Ssemwanga (2012) [92] | 88.3% | 90.0% | 0.0% | 73.3% | 95.0% |
| Wolf (2017) [93] | 99.8% | 99.8% | 61.1% | 99.3% | 98.2% |
| TenoRes Study Group (2016) [94] | 99.0% | 99.0% | 28.4% | 99.0% | 100.0% |
| van Zyl (2017) [95] | 94.9% | 93.2% | 57.6% | 93.2% | 94.9% |
| Huang (2003) [96] | 95.2% | 97.6% | 19.0% | 81.0% | 95.2% |
| **Average (unweighted)** | 94.3% | 93.3% | 34.1% | 84.2% | 78.9%<sup>2</sup> |
| **Average (weighted)** | 97.1% | 96.9% | 45.1% | 91.2% | 81.4%<sup>2</sup> |

<sup>1</sup> In this case, a substantial number of sequences that were classified as subtype A by REGA and our method were labeled unclassified subtypes (U) by COMET. In an HIV-1 phylogeny, subtype U sequences tend to be assigned a basal position (near the root) within the subtype A clade, suggesting that these sequences may be unrecognized variants or complex recombinants of subtype A.
<sup>2</sup> These low accuracies are primarily caused by REGA misclassifying many CRF01 sequences as subtype A, and subtype A is mostly equivalent to CRF01 in the *pol* region. If CRF01 and A were treated as equivalent, these accuracies would be 97.9% and 86.4% for the Rhee and Sukasem datasets, respectively, and unweighted and weighted averages of 93.8% and 96.2%, respectively.

The training set we use from the LANL 2010 web alignment does not have a balanced number of sequences for its 15 classes of HIV-1 subtypes or CRFs – classes range in size from 20 to 799 sequences. The smallest training set classes represented in our test data are subtype G and CRF 01B, each with 26 training sequences. To ensure that small training set classes do not negatively impact accuracy, we compute accuracy on those classes alone, and find a classification accuracy of 95.2% for subtype G and 100% for 01B, which is comparable to our accuracy on the rest of the test dataset.

Running time is another important performance indicator, so we also compare the performance of these five tools for the dataset of van Zyl et al. [95] and also for all datasets together (see Table 4). We observe that our tool matches or outperforms the competing state-of-the-art. Note that, for these comparison experiments, CASTOR, COMET, SCUEAL, and REGA were run from their web-based interfaces, and therefore the exact specifications of the machines running each programs could not be determined. For this reason, the running times presented here should be taken as rough order-of-magnitude estimates only.

**Table 4.** Approximate running times for all tested subtyping tools, for the dataset of van Zyl et al. [95] and all datasets listed in Table 3. The van Zyl dataset was chosen at random for this purpose.

| Tool | Running time for the van Zyl dataset | Running time for datasets from Table 3 |
| --- | --- | --- |
| KAMERIS | less than 2 seconds | 16 seconds |
| COMET | less than 2 seconds | 14 seconds |
| CASTOR | 3 seconds | 46 seconds |
| SCUEAL<sup>3</sup> | 18 minutes | 8 hours |
| REGA<sup>3</sup> | 31 minutes | 19 hours |

<sup>3</sup> The REGA and SCUEAL web servers have limits of 1000 and 500 sequences per run, respectively. Thus, 3 batches of sequences were needed for REGA, and 6 batches for SCUEAL to classify all sequences. COMET, CASTOR, and our tool have no such limits.

Overall, these experiments demonstrate our method is nearly identical in both accuracy and running time to the top third-party tool, COMET. Our tool differs from COMET in that it is open-source and freely available for commercial use, and is available in a standalone application which can be run on any computer, while COMET is closed-source and freely available for non-commercial research use only, and is publicly available only in a web-based system.

So far, we have only discussed supervised classification, and we have presented promising results for our approach. However, supervised classification requires data with known labels, which can be problematic considering that the rapid rates of mutation and recombination of viruses (particularly HIV-1) can lead to novel strains and recombinant forms emerging quickly. Unsupervised data exploration tools can help address this problem. To demonstrate, we take the set of all whole genomes from the LANL database and produce a MoDMap, visualizing their interrelationships, based on the Manhattan distance matrix obtained by computing all pairs of *k*-mer frequency vectors (see Methods section), for 9 different pure subtypes or groups (Fig 2), and just subtypes A, B, and C (Fig 3). As can be seen, based on these distances, the points in the plots are grouped according to known subtypes, and indeed it can be seen that subtypes A1 and A6 group together, and as well B and D group together, as could be expected.

**Fig 2.**
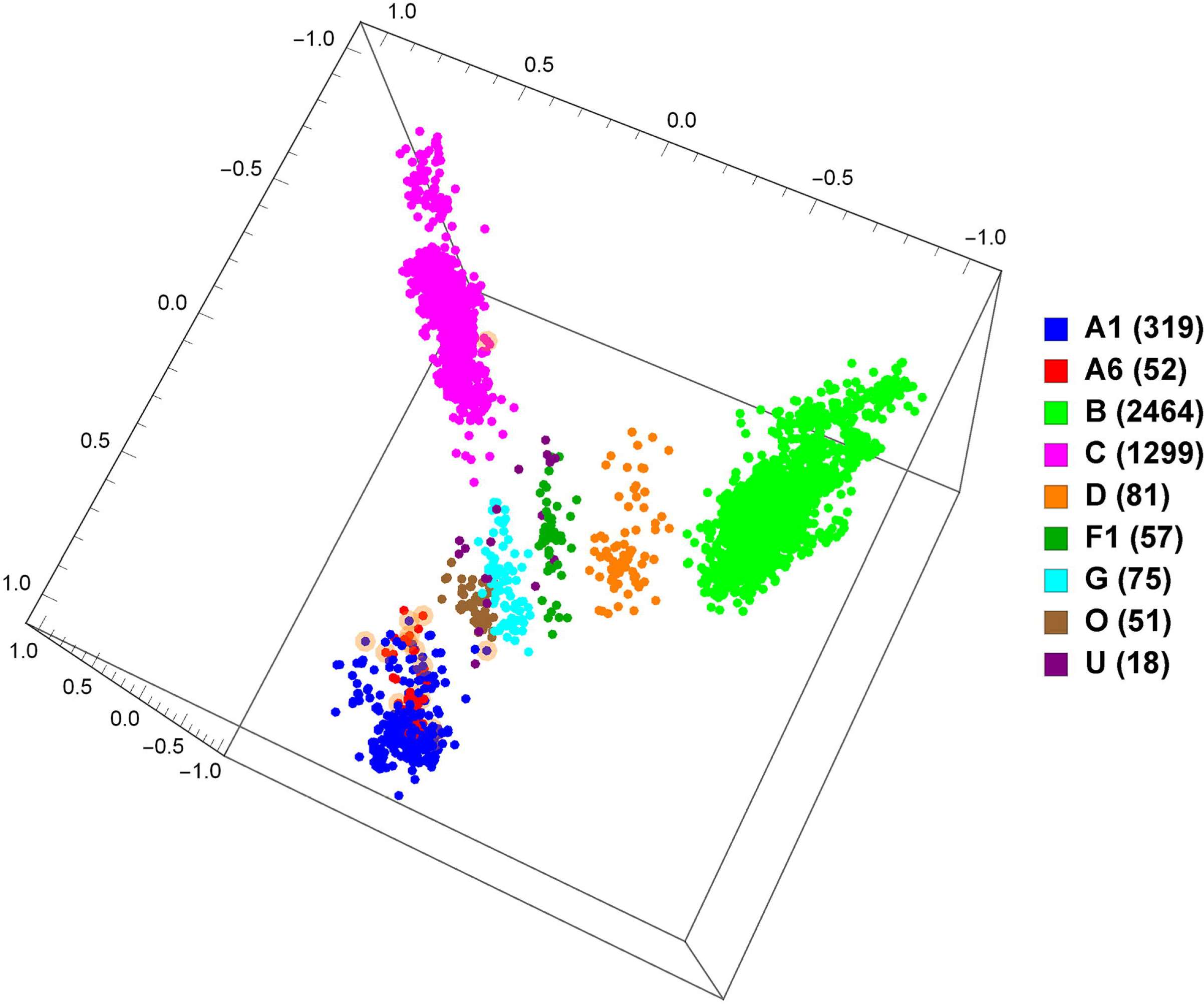
MoDMap of 4373 full-length HIV-1 genomes of 9 different pure subtypes or groups, at *k* = 6.

**Fig 3.**
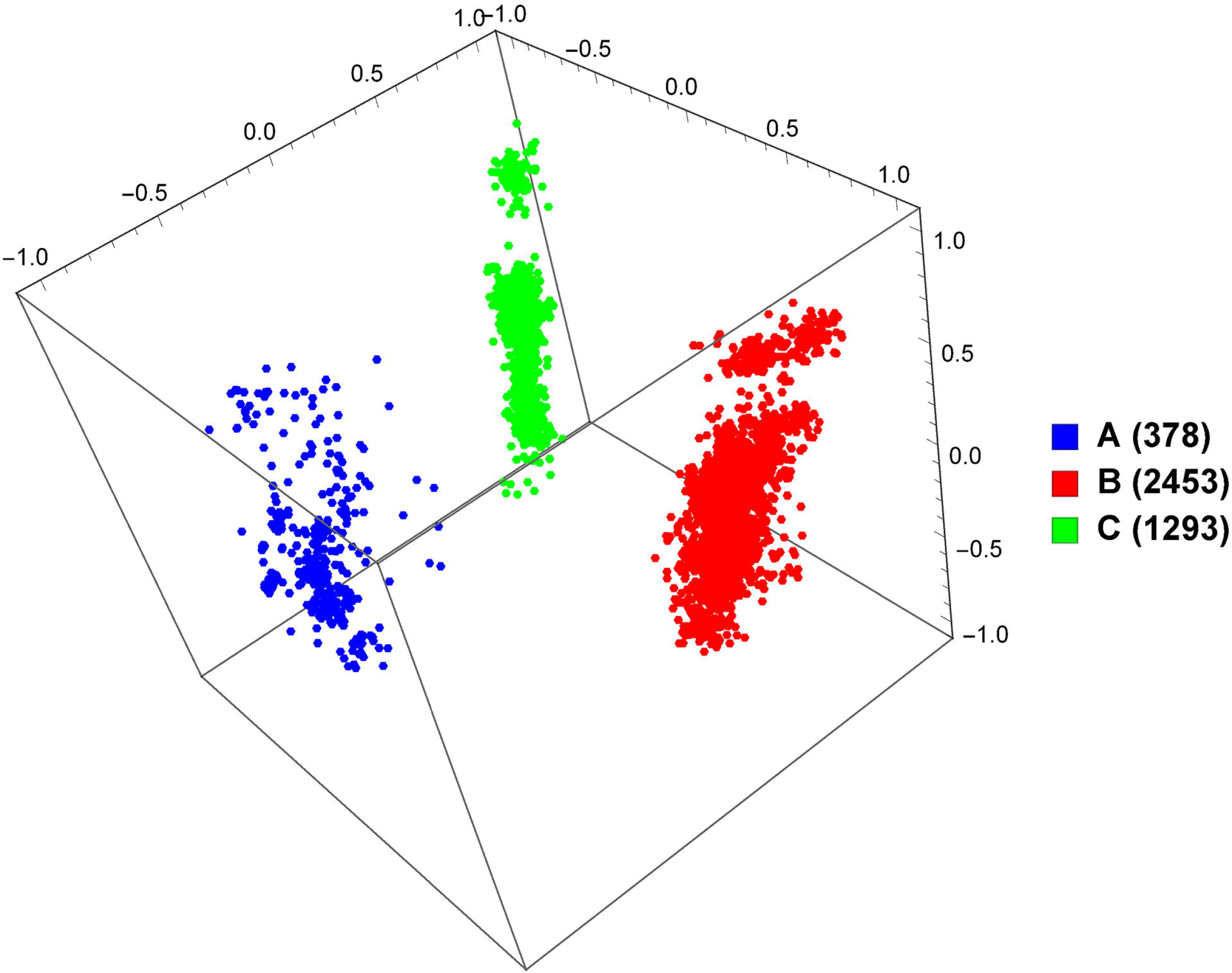
MoDMap of 4124 full-length HIV-1 genomes of subtypes A, B, and C, at *k* = 6.

Synthetic data has been useful in the study of viral species such as HIV-1, because a ground-truth classification is known for synthetic sequences without ambiguity. However, one may wonder how well such synthetic sequences model natural ones. We attempt to measure this by training a classifier on natural and synthetic HIV-1 sequence data – if natural and synthetic sequences cannot be distinguished, one may conclude that the simulation is realistic. For the ‘natural’ class we use the set of all *pol* genes from the LANL database, and for the ‘synthetic’ class we use 1500 synthetic *pol* genes produced as detailed above (see ‘Synthetic data’ in the Methods section), and we perform a 10-fold cross-validation at *k* = 6 and with the linear SVM classifier. We obtain an accuracy of 100%, meaning that the classifier can distinguish natural from synthetic sequences with perfect accuracy. This suggests that synthetic sequence data should be used with caution, since this result indicates it may not be perfectly representative of natural sequence data – specifically, our result suggests there is some characteristic of the synthetic sequences which differs from the natural sequences, which our method is able to recognize and use. We explore this further by generating a MoDMap, as seen in Fig 4. Interestingly, even though our supervised classifiers succeeded to discriminate between real and synthetic sequences with an accuracy of 100%, the approach using distances between *k*-mer frequency vectors results in the natural and synthetic sequences of specific subtypes grouping together, indicating that the synthetic sequences have some features that relate them to the corresponding natural sequences of the same subtype.

**Fig 4.**
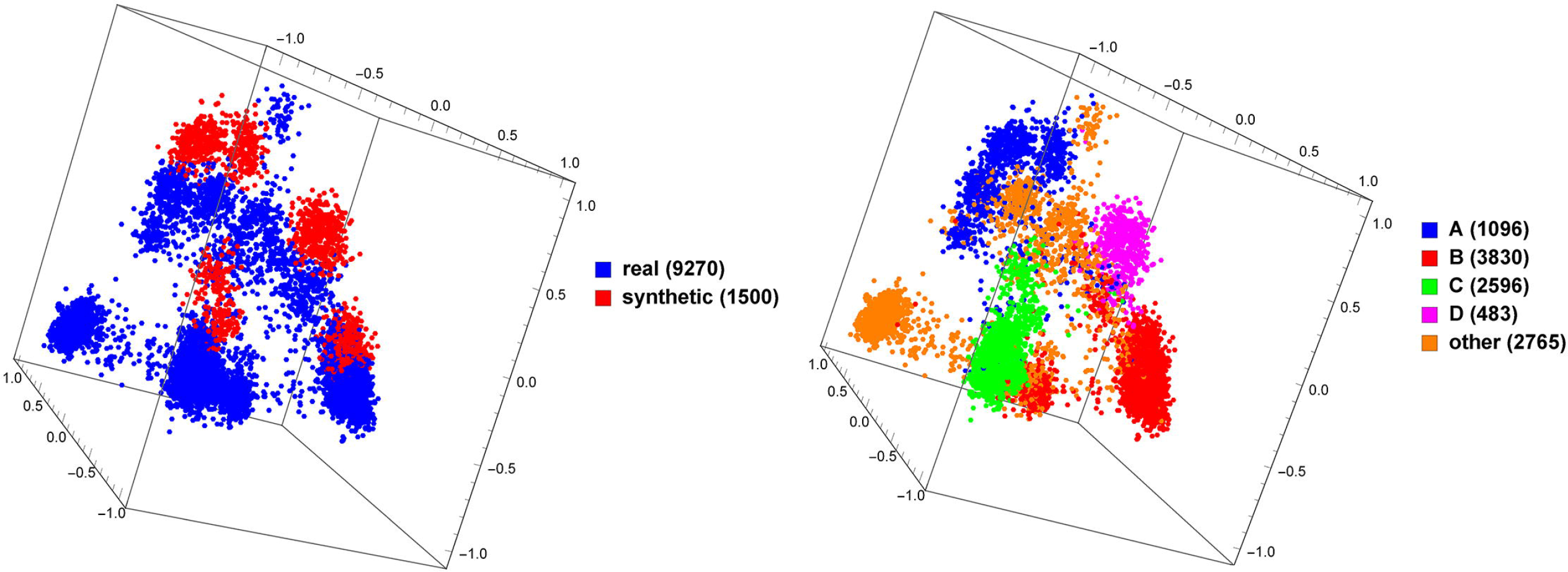
MoDMap of 9270 natural HIV-1 *pol* genes vs. 1500 synthetically generated HIV-1 *pol* genes of various subtypes. The same plot is colored on the left by type (natural and synthetic) and on the right by HIV-1 subtype.

## Discussion

The *k*-mer based supervised classification method we propose in this paper has several advantages compared to other popular software packages for the classification of virus subtypes. First, we have shown on several manually-curated data sets that *k*-mer classification can be highly successful for rapid and accurate HIV-1 subtyping relative to the current state-of-the-art. Furthermore, releasing our method as an open-source software project confers significant advantages with respect to data privacy, transparency and reproducibility. Other subtyping algorithms such as REGA [104] and COMET [8] are usually accessed through a web application, where HIV-1 sequence data is transmitted over the Internet to be processed on a remote server. This arrangement is convenient for end-users because there is no requirement for installing software other than a web browser. However, the act of transmitting HIV-1 sequence data over a network may present a risk to data privacy and patient confidentiality – concerns include web applications neglecting to use encryption protocols such as TLS, or servers becoming compromised by malicious actors. As a concrete example, the webserver hosting the first two major releases of the REGA subtyping algorithm [104] was recently compromised by an unauthorized user (last access attempt on November 27, 2017). In contrast, our implementation is available as a standalone program, without any need to transmit sequence data to an external server, eliminating those issues. In addition, our implementation is released under a permissive open-source license (MIT). In contrast, REGA [9] and COMET [8] are proprietary ‘closed-source’ software, making it impossible to determine exactly how subtype predictions are being generated from the input sequences.

Relying on a remote web server to process HIV-1 sequence data makes it difficult to determine which version of the software has been used to generate subtype classifications, and by extension difficult to guarantee that classification results can be reproduced. There is growing recognition that tracking the provenance (origin) of bioinformatic analytical outputs is a necessary component of clinical practice. For example, the College of American Pathologists recently amended laboratory guidelines on next-generation sequence (NGS) data processing to require that: “the specific version(s) of the bioinformatics pipeline for clinical testing using NGS data files are traceable for each patient report” [105]. In contrast to other tools, our standalone package makes it easy to use exactly the desired version of the software and thus enables precise reproducibility.

We now discuss some limitations of our approach. Like many machine learning approaches, our method does not provide an accessible explanation as to why a DNA sequence is classified a certain way, compared to a more traditional alignment-based method. In some sense, the classifiers act more as a black box, without providing a rationale for their results. Another issue is our requirement for a sizable, clean set of training data. As opposed to an alignment-based method that could function with even a single curated reference genome per class, machine learning requires several examples per training class, as discussed previously, to properly train. Finally, one issue common to any HIV-1 subtyping tool is the fact that recombination and rapid sequence divergence can make subtyping difficult, especially in cases where the recombinant form was not known at the time of training. Other tools are capable of giving a result of ‘no match’ to handle ambiguous cases, but our method always reports results from the classes used for training.

To more clearly demonstrate this last issue, we generate a random sequence of length 10,000 with equal occurrence probabilities for A, C, G, and T, and we ask the five subtyping tools evaluated in our study to predict its HIV-1 subtype. As expected, REGA gives a result of ‘unassigned’ and SCUEAL reports a failure to align with the reference. Our tool reports subtype ‘U’ with 100% confidence, CASTOR predicts HIV-1 group ‘O’ with 100% confidence, and COMET reports SIV_CPZ_ (simian immunodeficiency virus from chimpanzee) with 100% confidence. These outcomes are consistent with the disproportionately large genetic distances that separate HIV-1 group O and SIV_CPZ_ from HIV-1 group M – a line drawn from a random point in sequence space is more likely to intersect the branch relating either of these distant taxa to group M. Similarly, branches leading to subtype U sequences tend to be longer and to intersect the HIV-1 group M tree at a basal location^4^. This artificial example implies that real HIV-1 sequences that do not readily fit into any of the defined subtypes or circulating recombinant forms may result in incorrect predictions with misleadingly high confidence scores.

In spite of these limitations, our method not only matches or improves upon current HIV-1 subtyping algorithms, but it should also be broadly applicable to any DNA sequence classification problem, including other virus subtyping problems. To demonstrate this, we use the same method (with *k* set to 6 and a linear SVM classifier) and 10-fold cross-validation to measure the accuracies for classifying dengue, hepatitis B, hepatitis C, and influenza type A virus full-length genomes (described in the Datasets section) to their respective reference subtypes. Overall, we obtain accuracies of 100% for dengue virus, 95.81% for hepatitis B virus, 100% for hepatitis C virus, and 96.68% for influenza A virus. We also provide a MoDMap visualization of the subtypes of hepatitis B, as seen in Fig 5. This plot displays not only clear separation between subtypes but also structure within subtypes A and B, which would be an interesting target for future study.

**Fig 5.**
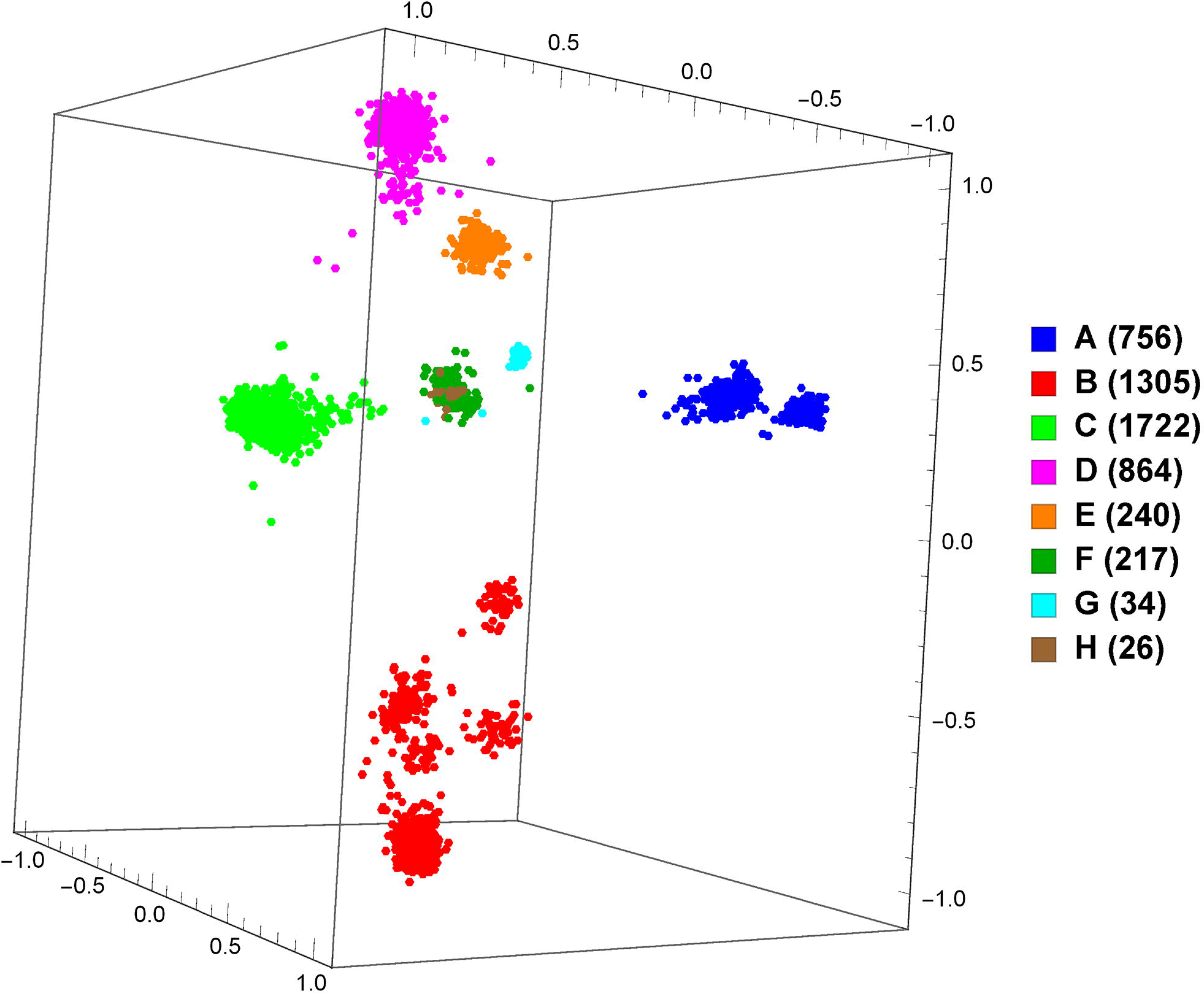
MoDMap of 5164 whole hepatitis B genomes of 6 different pure subtypes.

Because of the exponential growth of sequence databases, modern bioinformatics tools increasingly must be capable of handling NGS sequence data and must be scalable enough to manage huge sets of data. As well, researchers often demand the privacy, security, and reproducibility characteristics an open-source, standalone, offline tool such as ours provides. However, there remain several areas for future work. Although our tool matches or exceeds the classification speed of the competing state-of-the-art, performance optimization was not a focus of this study and we believe there is room to substantially improve running time even further. Similarly, although we match or exceed the classification accuracy of the competing state-of-the-art, different modern machine learning methods such as GeneVec [106] or deep neural networks may permit us to achieve even higher accuracy on challenging datasets. As well, given the rapid rate of mutation of many viruses, it would be highly useful for our tool to be capable of giving a result of ‘no match’ with its training data. Further, since our software is able to classify sequences quickly, we believe it should be capable of handling large NGS sequence datasets, and we leave a careful study of this application to future work. Each of these possibilities could make our method and software even more useful in the future.

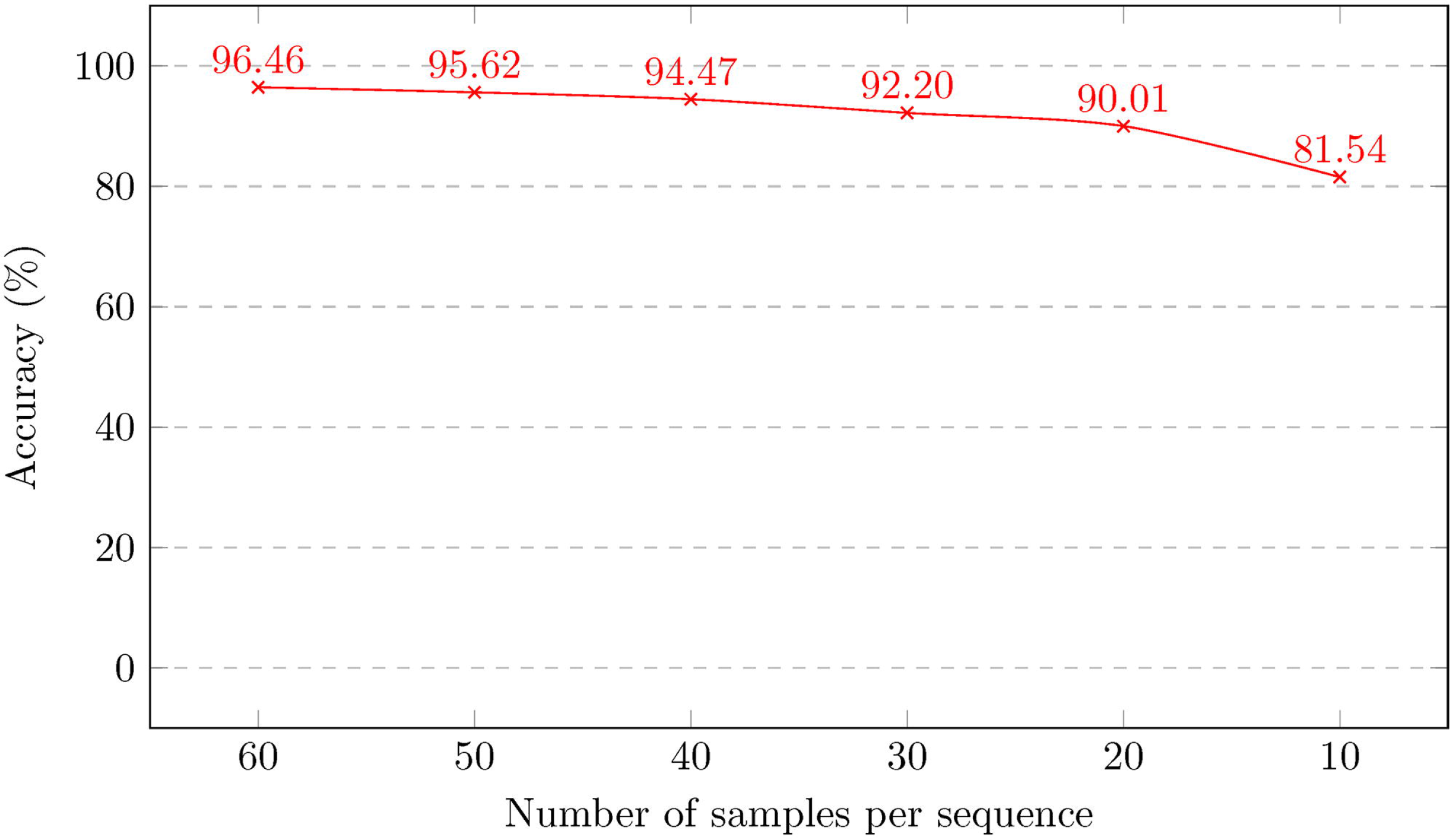

## Supporting information

**S1 Appendix Instructions for reproduction of the experiments using Kameris.**

**S2 Appendix List of subtypes of viral species from datasets used.**

## Acknowledgments

We wish to thank David Nguyen for his contributions to running preliminary datasets of simulated HIV sequence data sets through the web-based subtyping methods, Gurjit Randhawa for helpful technical and implementation-related discussions, and Dr. Stephen Watt for suggesting the name Kameris for this software.

## Footnotes

4 HIV-1 subtype U does not comprise a distinct clade. Rather, the LANL database labels sequences as ‘U’ when they belong to a lineage not meeting the criteria required for a designation as a subtype [5]. However, practical but anecdotal experience suggests subtype U sequences are typically basal.

